# Clonality, specificity, and HLA restriction of human T cell receptors against *Plasmodium falciparum* CSP

**DOI:** 10.1101/2025.01.06.631573

**Authors:** Hannah van Dijk, Ilka Wahl, Sara Kraker, Paul M. Robben, Sheetij Dutta, Hedda Wardemann

**Affiliations:** B Cell Immunology, German Cancer Research Center, Heidelberg, Germany; Biologics Research and Development Branch, Center for Infectious Diseases Research, Walter Reed Army Institute of Research, Silver Spring, MD, United States

## Abstract

*Plasmodium falciparum* malaria remains a significant global health challenge. Current vaccines elicit antibody responses against circumsporozoite protein (PfCSP) that prevent the infection of hepatocytes but offer only moderate protection. Cellular immunity has emerged as a critical component of pre-erythrocytic protection that might be leveraged to develop improved PfCSP vaccines. Here, we characterized the clonality, molecular features, epitope specificity, and HLA-restrictions of the human PfCSP-specific CD4^+^ and CD8^+^ T cell response to vaccination with an adjuvanted PfCSP vaccine, FMP013/ALFQ. Using TCR expression cloning, we identified novel conserved CD4^+^ T cell epitopes in the PfCSP N terminus and show that the C-terminal CS.T3 epitope was targeted by CD4^+^ and rare CD8^+^ T cells, which recognized this epitope co-receptor-independently presented on a class II HLA. Our findings provide insights into the utility of these epitopes as targets for strain-transcending immunity compared to the immunodominant but highly polymorphic epitopes in the PfCSP C terminus, offering guidance for the design of improved malaria vaccines.

## Introduction

Malaria remains a significant global health issue, with *Plasmodium falciparum* (*Pf*) responsible for the most severe cases and highest mortality rates. The only available vaccines, RTS,S/AS01 and R21/MM, target *Pf* circumsporozoite protein (PfCSP), the major surface antigen of *Pf* sporozoites. Both vaccines rely primarily on the induction of anti-PfCSP antibodies that block sporozoites during their brief migration from the site of inoculation in the skin to the liver, preventing hepatocyte invasion and subsequent progression to the erythrocytic stage of infection, which causes disease (Agnandji et al., 2011; RTS.S Clinical Trials Partnership, 2014; Casares et al., 2010; Datoo et al., 2022; Datoo et al., 2024). Despite inducing strong antibody responses, these vaccines provide protection, which wanes rapidly over time, particularly in high-transmission regions (Bejon et al., 2013).

T cells likely contribute to RTS,S/AS01-induced protection by providing B cell help and promoting antibody affinity maturation as suggested by the finding that PfCSP-specific CD4^+^ T follicular helper (Tfh) cell responses were associated with improved protection (Pallikkuth et al., 2020). However, the role of T cells in pre-erythrocytic immunity against *Pf* goes beyond complementing humoral responses. Accumulating evidence from vaccination studies with radiation- or genetically-attenuated sporozoites suggests that cellular immunity including PfCSP-specific T cells critically contributes to protection (Hassert et al., 2023; Oliveira et al., 2008; Rodrigues et al., 1991; Romero et al., 1989; Seder et al., 2013). In particular, CD4^+^ T cells might play a strong and multifaceted role through the production of IFN-γ, which stimulates class I and II-mediated antigen presentation, promotes the activation of cytotoxic T and NK cells, and mediates suppression of parasite growth in infected host cells, as well as through their direct cytolytic activity (Rénia et al., 1993; Rénia et al., 1991; Weiss et al., 1993; Romero et al., 1989). Indeed, clinical immunity induced by natural parasite exposure seems to be linked to the clonal expansion of CD4^+^ T cells with a cytotoxic phenotype against pre-erythrocytic antigens including PfCSP (Nardin et al., 1989; Moreno et al., 1991; Furtado et al., 2023).

How CD8^+^ T cells against PfCSP or other *Pf* antigens contribute to malaria immunity in humans is poorly understood (Good et al., 1987; Moreno et al., 1991; Good et al., 1988; Sinigaglia et al., 1988b; Moreno et al., 1993; Sinigaglia et al., 1988a; Doolan et al., 1997; Heide et al., 2019), despite strong preclinical evidence based on adoptive T cell transfer and depletion experiments as well as studies with T cell-deficient animals showing that CD8^+^ T cells confer protection by eliminating infected hepatocytes via perforin-mediated lysis and IFN-γ secretion (Romero et al., 1989; Rodrigues et al., 1991; Weiss et al., 1988; Schofield et al., 1987). The assessment of CD8^+^ T cell responses to *Pf* in humans is hampered by the relative weakness and fast decline of CD8^+^ responses in circulation after vaccination and their tissue residency at the site of infection, i.e. the liver after natural or immunization-mediated exposure to sporozoites (Weiss and Jiang, 2012; Trimnell et al., 2009; Schofield et al., 1987; Hoffman et al., 1989).

Despite the evidence that PfCSP-specific CD4^+^ and potentially also CD8^+^ T cells play a role in protection from malaria, little is known about the molecular characteristics of TCRs that mediate PfCSP binding to inform the design of vaccines with the potential to mediate more potent protection. PfCSP consists of a central repeat region flanked by a short junctional peptide, which connects the repeat to the N-terminal domain, and a C-terminal domain. Bulk cell analyses have shown that T cell epitopes are distributed across the junctional peptide and N- and C-terminal domains, but are absent from the central repeat region. CD4^+^ T cells mainly target Th2R and T*, two epitopes in the C-terminal α-thrombospondin type-I repeat (αTSR) domain, with weaker responses against CS.T3 in the C terminus and T1 in the N-terminal junction (Moreno et al., 1993; Moreno et al., 1991; Wahl et al., 2022b; Good et al., 1987). Differences in immunogenicity among these epitopes are linked to HLA restrictions and differences in sequence conservation of the target peptides with stronger immune responses to the polymorphic epitopes Th2R and T* that are presented on diverse HLA alleles, compared to the more conserved CS.T3 and T1 epitopes with stronger HLA restrictions (Good et al., 1988; Schwenk et al., 2011).

The polymorphic nature of the immunodominant C-terminal epitopes poses a challenge to vaccine development, as it limits cross-strain immunity (Zeeshan et al., 2012; de la Cruz et al., 1988; Neafsey et al., 2015). A detailed understanding of epitope-specific T cell responses, including their HLA restriction and TCR usage, is critical to overcome these limitations and guiding the design of next-generation malaria vaccines that elicit robust, strain-transcending T cell and antibody responses for broad and efficacious protection (Beura et al., 2018).

We have recently characterized the PfCSP-specific circulating T follicular helper cell response to irradiated sporozoite vaccination at single-cell level, which identified highly strain-specific TCRs against supertopes in the polymorphic Th2R/T* region (Wahl et al., 2022b). In this study, we analyzed the peripheral PfCSP-specific CD4^+^ and CD8^+^ T cell responses in malaria-naive individuals vaccinated with FMP013, a nearly full-length recombinant PfCSP adjuvanted in Army Liposome Formulation containing QS21 (ALFQ), which induced strong antibody responses against the C terminus and gradually weaker responses against the central repeat and N-terminal domains in a recent Phase I study (Hutter et al., 2022). Using single-cell TCR sequencing, expression cloning, and functional assays, we identified novel and conserved epitopes within the N- and C-terminal domains. We further characterized the gene usage, HLA restriction, and functional properties of these TCRs, revealing a highly polyclonal, donor-specific CD4^+^ T cell repertoire dominated by responses to C-CSP. Our findings also provide new insights into the rare CD8^+^ T cell responses against PfCSP, including a co-receptor-independent TCR targeting a shared CD4^+^/CD8^+^ epitope presented in an HLA-DP-restricted manner. These data expand our understanding of PfCSP-specific T cell immunity and highlight CS.T3 as an important target for the development of broadly protective malaria vaccines.

## Results and discussion

### CD4^+^ T cell response to repeated PfCSP vaccination

To characterize the human CD4^+^ T cell response to PfCSP, we isolated peripheral blood mononuclear cells (PBMCs) from seven malaria-naive individuals (F1-5, F7, F9) isolated four weeks after vaccination with two doses of recombinant Pf3D7 strain-based PfCSP vaccine FMP013 in the Army Liposome Formulation containing QS21 (ALFQ), an adjuvant that also contains the synthetic monophosphoryl lipid A derivative analog 3D-PHAD^®^ (Figure 1A; (Hutter et al., 2022). To identify PfCSP-reactive T cells, we first stimulated PBMC samples from two donors (F7, F9) with pools of overlapping peptides covering the complete immunogen PfCSP amino acid (aa) sequence (3D7 aa Tyr_26_-Ser_383_; Figure 1B). Flow cytometric analyses showed an increase in the number of activated CD4^+^ T cells expressing the activation markers CD25 and OX40 compared to unstimulated control cultures from both donors demonstrating that antigen-reactive CD4^+^ T cells were circulating in blood 28 days after the second FMP013/ALFQ vaccination. To determine the clonal composition of the activated T cells, we isolated single CD4^+^CD25^+^OX40^+^ T cells from the stimulated cultures by indexed flow cytometric cell sorting and amplified and sequenced their paired *TRA* and *TRB* genes (Supp Figure 1A). Only about half of the sequences (47% in F7, 53% in F9) were obtained from lowly expanded clones encoded by diverse V segments reflecting the highly polyclonal nature of the anti-FMP013 response in both individuals (Figure 1C).

**Figure 1:**
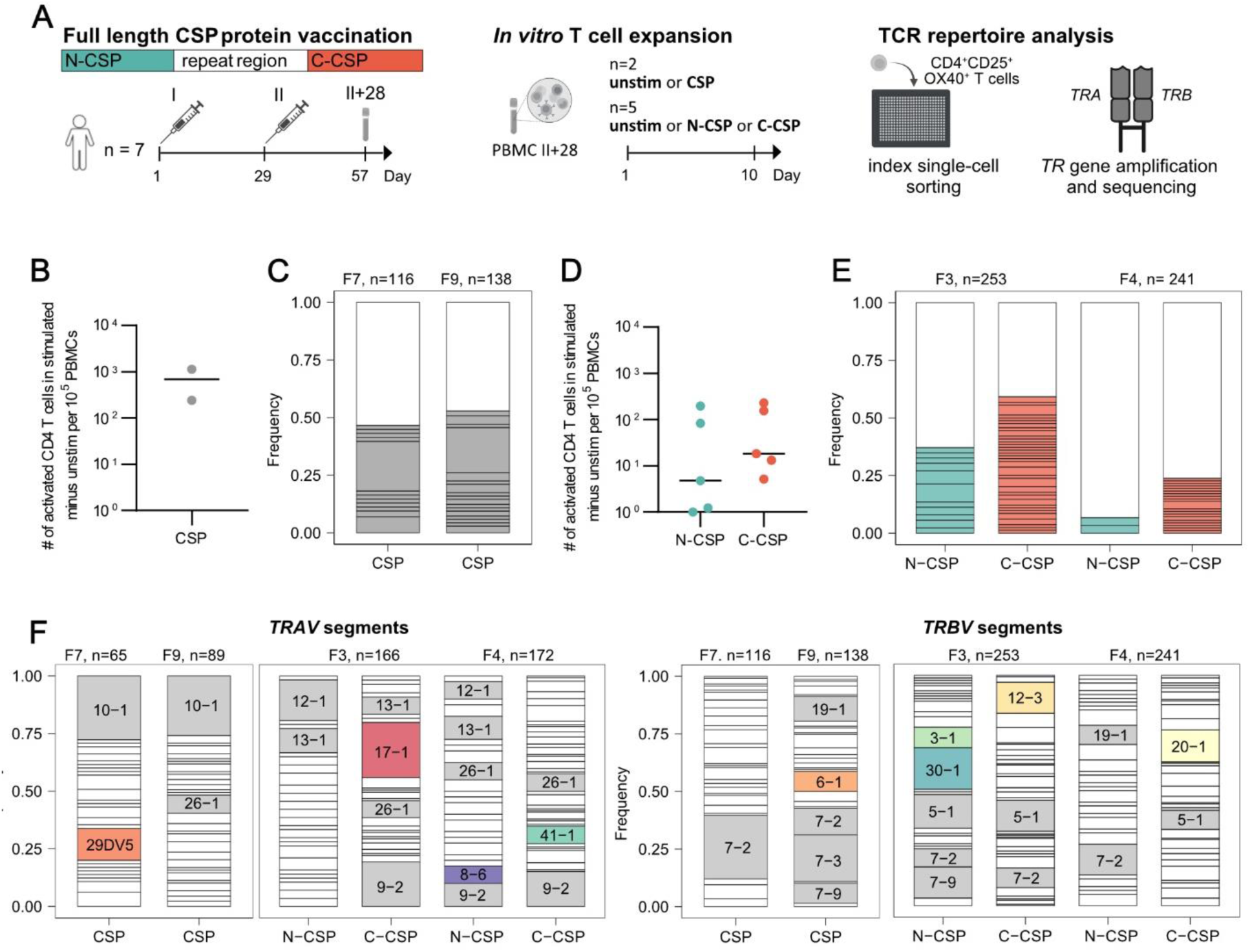
Vaccination induced polyclonal CD4^+^ T cells targeting the PfCSP N and C terminus. **A** Schematic overview of the sample collection, *in vitro* stimulation, and single cell isolation strategy. PBMC samples collected 28 days after two FMP013/ALFQ vaccine dose (II+28) were left untreated or stimulated with peptide pools covering the complete FMP013 aa sequence (CSP), the FMP013 N-terminus, junction, and repeat domain (N-CSP), or the FMP013 C-terminus (C-CSP). Indexed flow-cytometric single-cell sorting was performed ten days later. **B and D** Number of activated (CD25^+^OX40^+^) CD4^+^ T cells after *in vitro* peptide pool stimulation compared to unstimulated control cells from the same individuals. Each symbol represents a volunteer. **C and E** Clonal composition and *TRAV* and *TRBV* gene segment usage of single activated CD4^+^ T cells after CSP (C) or N-CSP and C-CSP (E) peptide pool stimulation of samples from the indicated donors. Individual expanded clones are shown in color, all non-expanded clones are shown in white. **F** *TRAV* and *TRBV* gene usage after *in vitro* stimulation, *TRAV* and *TRBV* gene segments (>7%) enriched in activated cells isolated from peptide-stimulated cultures are labelled and non-overlapping segments highlighted in color.

To distinguish the response to the individual PfCSP subdomains, we stimulated PBMCs from the other five donors with peptide pools covering either the N terminus, junction region, and repeat domain (N-CSP) or the complete C terminus (C-CSP; Figure 1D). T cells from all individuals responded to stimulation with C-CSP peptides, whereas cultures from only three of the five donors responded to N-CSP peptide stimulation. TCR gene sequencing of the activated cells from the two donors with the strongest N- and C-CSP responses showed that the degree of clonal expansion was lower after N-CSP (36% in F3, 7% in F4) compared to C-CSP peptide pool stimulation (61% in F3, 27% in F4), in line with previous reports demonstrating the immunodominance of C-terminal compared to N-terminal and junction epitopes (González et al., 2000)). The absence of any clonal overlap between the cultures and differences in *TRAV* and *TRBV* gene enrichment after N- and C-CSP peptide pool stimulation and between the donors indicated that the cells expanded peptide and HLA specifically (Figure F; *TRAV29-DV5, 17-1, 8-6, 41-1; TRBV30-1, 3-1, 12-3, 20-1, 6-1*). In summary, FMP013/ALFQ vaccination induced a highly polyclonal, donor-specific CD4^+^ T cell response against the PfCSP C terminus in all donors, whereas responses against the N terminus, junction and repeat domain were induced in only a few individuals.

### Polyclonal PfCSP-specific CD4^+^ T cells target N-terminal, junction, and C-terminal epitopes

To determine whether the activated and expanded T cell clones were PfCSP-reactive, we cloned the TCRs of 66 selected clones, including several with enriched gene segments after the stimulation, and expressed them in CD4^+^ positive TCR-deficient Jurkat76 cells (Supp Figure 1B, Supp Table 1). More than half (53%, 35/66) of the T cell lines secreted IL-2 after *in vitro* co-culture with autologous B cells pulsed with the C-CSP or N-CSP peptide pools (Figure 2A). Two thirds (69%, 24/35) showed specificity for C-CSP peptides, whereas one third (31%, 11/35) reacted exclusively to N-CSP peptides (Figure 2B). Stimulation of these clones with individual peptides identified their target epitopes (Figure 2C, D). The N-CSP specific TCRs reacted to two novel highly conserved epitopes covering aa 61-75 (4/11) and 81-95 (1/11), hereafter referred to as CSP61 and CSP81, respectively, or to the T1 epitope in the N-terminal junction (6/11) but not to the repeat (Figure 2C, D). The C-CSP-reactive TCRs recognized peptides covering the known polymorphic Th2R (peptide 315-319;15/24) and T* epitopes (peptide 319-337; 2/24) as well as CS.T3 (peptide 367-381; 7/24; Figure 2C; (Sinigaglia et al., 1988b; Good et al., 1988; Guttinger et al., 1988). Although the PfCSP-reactive TCRs were encoded by diverse *TRAV* and *TRBV* combinations, we observed several associations between epitope specificity and V gene usage (Figure 2E, F). Specifically, T1 binding was linked to *TRBV30-1* usage as previously reported (Wahl et al., 2022b), whereas the majority of Th2R and CS.T3 reactive TCRs were encoded by *TRAV17-1* and *TRBV20-1*, respectively. In summary, we identified CD4^+^ T cell-associated TCRs against epitopes in the PfCSP C- and N terminus, including two novel N-terminal epitopes and defined TCR gene usage patterns associated with reactivity against several of these targets.

**Figure 2:**
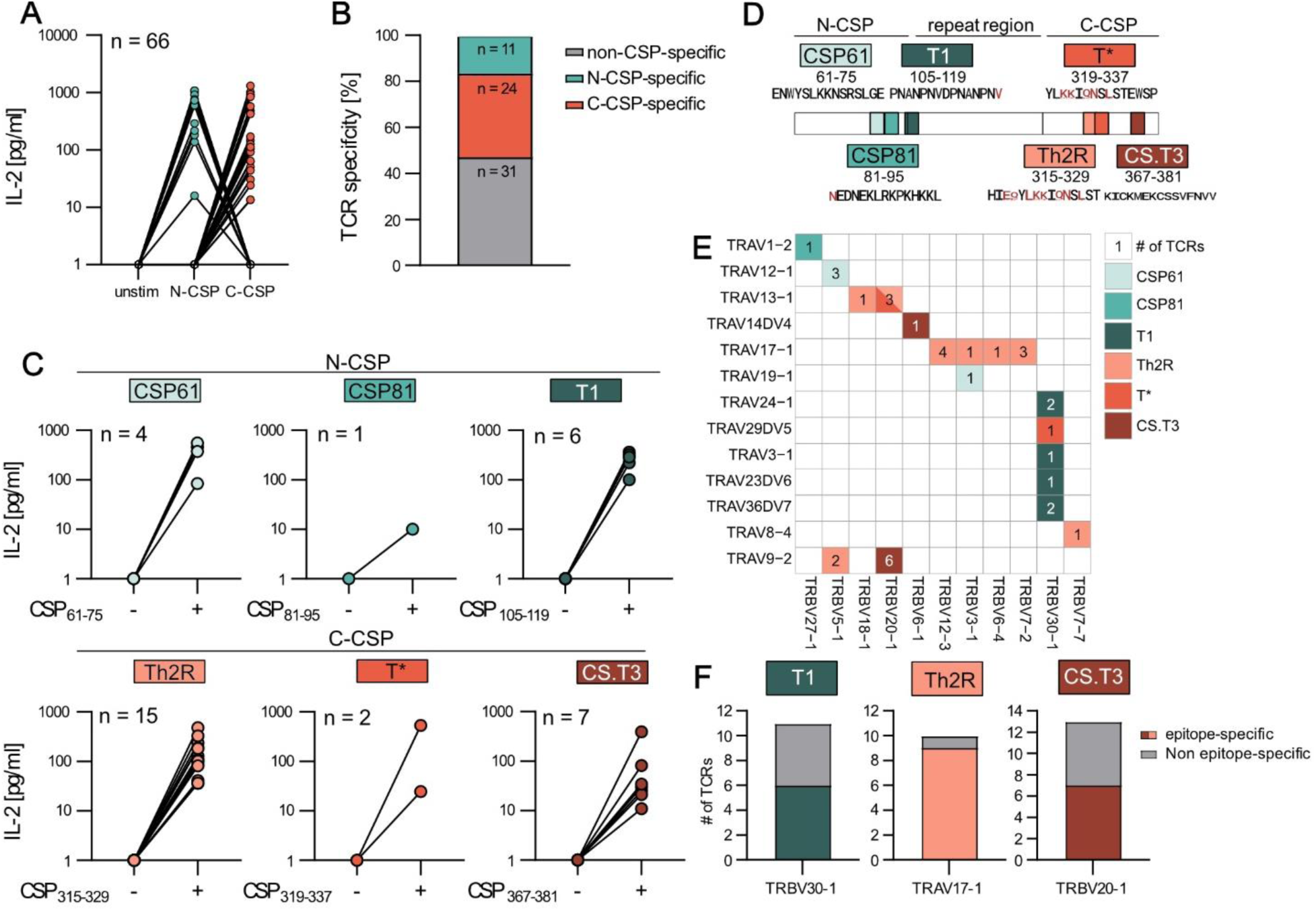
PfCSP-specific CD4^+^ T cells target three N-CSP and three C-CSP epitopes. Transgenic CD4^+^ Jurkat76 T cell lines expressing TCRs from T cells with an activated phenotype after stimulation of PBMCs with N- or C-CSP peptide pools (donors F3, F4, F7, F9) were generated. **A** IL-2 concentrations as quantified by ELISA in supernatants of T cell lines (n=66) co-cultured with autologous B cells pulsed with N-CSP or C-CSP peptide pools or with non-peptide pulsed autologous B cells (unstim). **B** Frequency of N- and C-CSP-reactive and non-reactive TCRs. **C** IL-2 concentrations as quantified by ELISA in supernatants of T cell lines co-cultured with autologous B cells pulsed with the indicated peptides (+) or left unstimulated (−). n indicates the number of tested T cell lines. **D** Schematic map of the identified target epitopes and aa position within PfCSP. Sequence diversity of TCR epitopes among 481 PfCSP sequences isolated from 7 geographic regions published by (Tanabe et al., 2013), polymorphic aas are highlighted in red. **E** *TRAV* and *TRBV* usage and number of individual TCRs with the indicated epitope specificity. **F** Number of epitope-specific TCRs among all cloned and tested TCRs with the indicated V segments. **A, C** One out of two independent experiments is shown.

### PfCSP-specific TCRs are restricted to HLA alleles with variable prevalence

To determine the HLA restriction of the peptide-specific TCRs, we performed the co-culture stimulation assay with autologous B cells in the presence of HLA-DR, DQ, and DP blocking antibodies (Figure 3A). The anti-HLA-DR antibody exclusively blocked the activation of all T cell lines expressing CSP61-, CSP81-, Th2R- and CS.T3-specific TCRs, whereas the activation of T1-specific T cell lines was blocked specifically by anti-HLA-DQ. In contrast, activation of the T*-specific TCRs was strongly inhibited by HLA-DR or HLA-DQ blocking antibodies.

**Figure 3:**
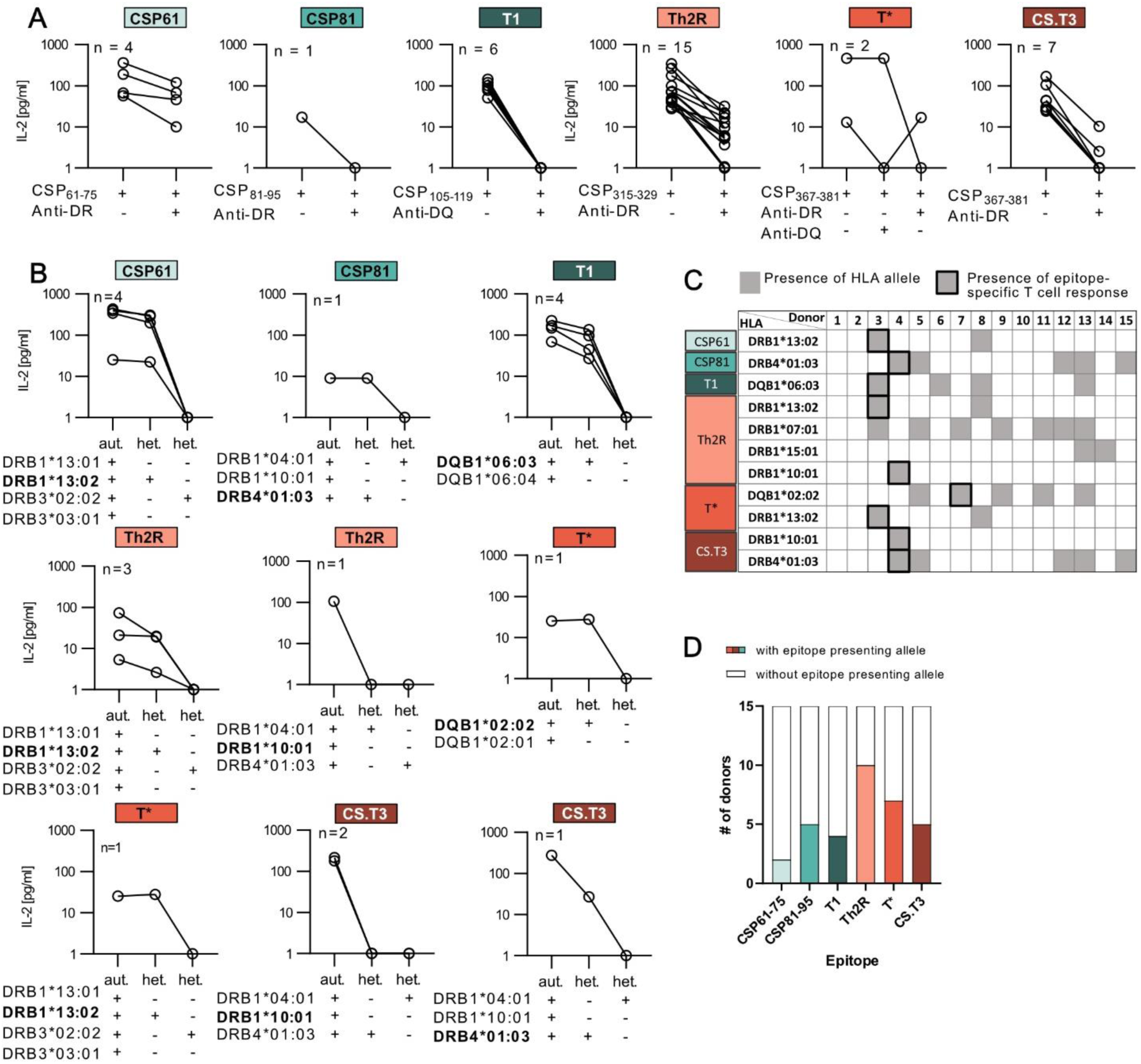
PfCSP epitopes are presented on HLA alleles with variable prevalence. **A** IL-2 concentrations as measured by ELISA in supernatants of TCR-transgenic CD4^+^ Jurkat76 T cells co-cultured with autologous B cells pulsed with the indicated peptides in the presence (+) or absence (−) of the indicated locus-specific HLA-blocking antibodies. **B** IL-2 concentrations as measured by ELISA in supernatant of TCR-transgenic CD4^+^ Jurkat76 T cells co-cultured with peptide-pulsed autologous (aut.) or heterologous (het.) B cells that share one (+) or no (−) locus-specific HLA allele with the respective autologous B cells. Epitope-presenting HLA alleles were determined (activation observed in co-cultures with heterologous B cells expressing a single share allele) or inferred (lack of activation in heterologous B cells with one shared allele) and are highlighted in bold. **C** HLA-alleles presenting the indicated PfCSP epitopes in the 15 donors described in this study (DRB1*13:02, DRB1*04:03, DQB1*06:03, DRB1*10:01, DQB1*02:02) or previously (DRB1*07:01, DRB1*15:01 (Wahl et al., 2022b)). Gray shading indicates the presence of an allele. Black frames indicate epitope-specific CD4^+^ T cell responses as determined in (A). **D** Frequency of donors with HLA alleles shown to present the indicated epitopes. **A, B** n indicates the number of tested T cell lines. One out of two independent experiments is shown.

Next, to define or infer the HLA allele specificity of all TCRs, we performed heterologous stimulation experiments of the T cell lines with peptide-pulsed B cells from donors that shared only a single or no HLA-allele (Figure 3B). All TCRs that targeted N-terminal epitopes (CSP61, CSP81, T1) recognized their specific target peptide in the context of a single HLA allele (DRB1*13:02, DRB4*01:03 and DQB1*06:03, respectively). In contrast, the TCRs with reactivity against the C terminus were restricted to two alleles. The two Th2R-specific TCRs were activated by peptides presented in the context of DRB1*13:02 and DRB1*10:01, whereas the two T*-specific TCRs were restricted to DQB1*02:02 and DRB1*13:02, and the CS.T3-specific TCRs recognized their epitope on DRB1*10:01 and DRB4*01:03. The number of individuals that expressed the above alleles or alleles that we previously reported to present Th2R peptides (DRB1*07:01 or DRB1*15:01 (Wahl et al., 2022b)) varied strongly among the donors analyzed in this study (Figure 3C). The majority of donors expressed HLA alleles that induced T cell responses against the polymorphic Th2R and T* epitopes (10/15 and 7/15, respectively), reflecting the high abundance of TCRs with specificity for peptides covering this region (Figure 3D). Alleles presenting the conserved CS.T3 and N-terminal peptides were less frequent in line with the lower frequency of TCRs with these specificities (Figure 2). In summary, we identified novel and abundant HLA alleles presenting the immunodominant Th2R and T* peptides and determined the allele context for TCRs against novel conserved CD4^+^ T cell epitopes in the PfCSP N terminus (CSP61, CSP81) and the conserved CS.T3 epitope in the C terminus.

### N- and C-terminal PfCSP peptides are presented on diverse HLA class I alleles

Several of the CD4^+^ T cell epitopes including the newly identified conserved CSP61 and CSP81 epitopes and CS.T3 have previously been associated with HLA class I-restricted CD8^+^ T cell responses (Wang et al., 1998; González et al., 2000; Doolan et al., 1991; Sedegah et al., 2013), which prompted us to investigate the CD8^+^ T cell response against PfCSP in the same vaccinated individuals. We first used NetMHCIpan4.0 (Reynisson et al., 2020), a computational peptide-MHC prediction algorithm, to estimate the binding strength of the different HLA class I molecules expressed by the donors in our study to peptides covering the complete FMP013 aa sequence (Figure 4A). Seven peptides, two N-terminal peptides (CSP_65-73_, CSP_86-94_) and five C-terminal peptides (CSP_285-293_, CSP_310-319_, CSP_319-327_, CSP _336-345_, CSP_353-360_) were predicted to bind strongly to three or more of the 13 HLA-A and B alleles that were detected in our cohort. For five HLA alleles with high-prevalence (HLA-A*01:01, HLA-A*02:01, HLA-A*11:01) or high number of strong predicted binders (HLA-B*07:02, HLA-B*08:01), peptide binding was confirmed using HLA monomers loaded with the predicted N-terminal or C-terminal peptides (Figure 4B). In line with the binding predictions, HLA-B*07:02 and HLA*B08:01 bound more peptides with higher apparent affinity than the more frequent HLA-A alleles A*01:01, A*02:01, and A*11:01 (Figure 4C). The detection of HLA-A and B alleles that can present N- and C-terminal peptides prompted us to assess whether we could detect anti-PfCSP CD8^+^ responses.

**Figure 4:**
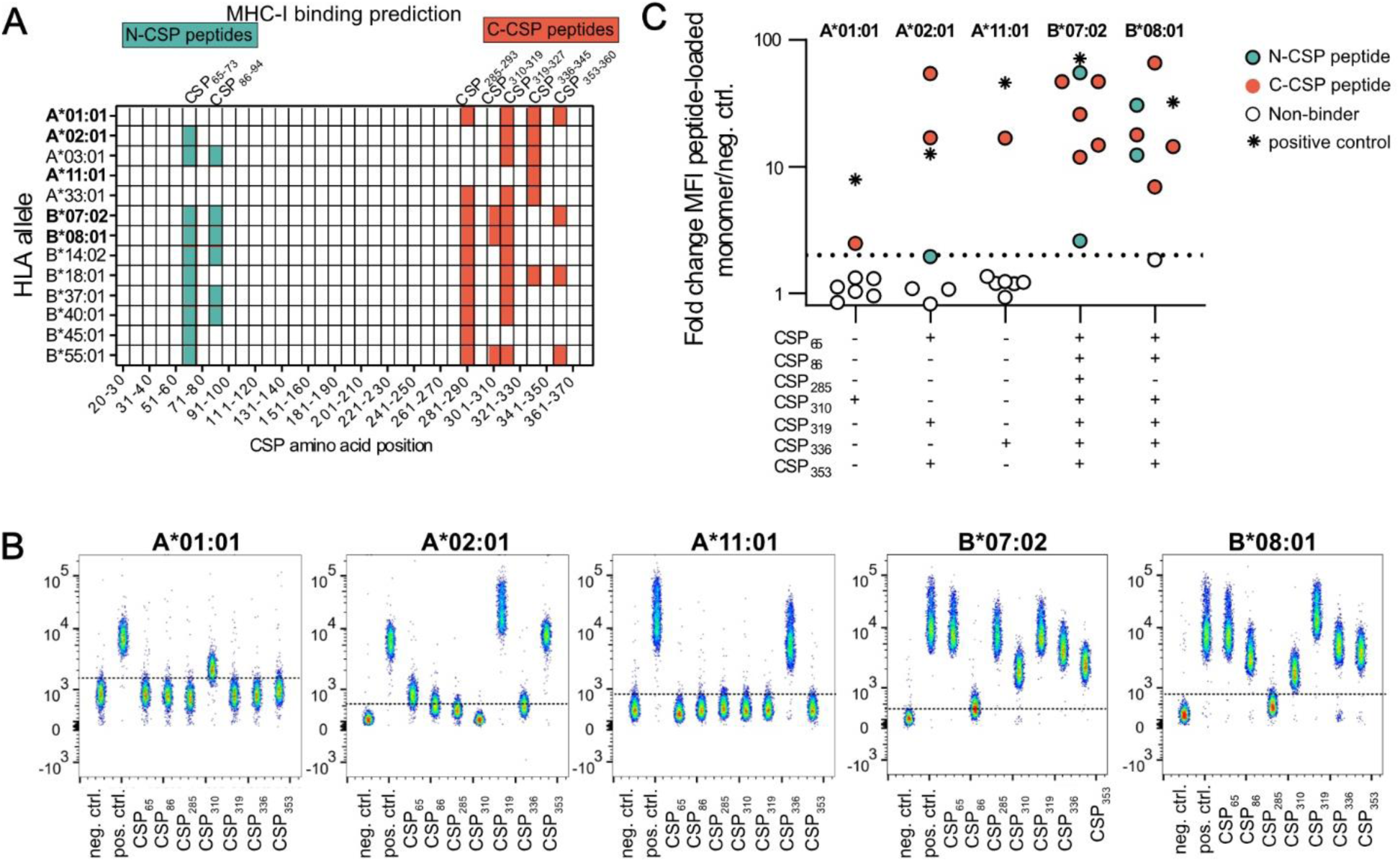
N- and C-terminal peptides bind HLA class I alleles promiscuously. **A** Peptide binding prediction of PfCSP peptides to HLA-A and HLA-B alleles of five vaccinated donors by NetMHCpan4.1 tool. Predicted peptides (9-11mer length, EL-rank threshold below 7) within the FMP013 aa sequence (position 20-380) are highlighted in color and shown within starting position ranges of 10 aa. **B-C** Binding of 7 predicted PfCSP peptides to five HLA monomer. **B** FACS plot of ß-microglobulin stained monomers loaded with individual peptides compared to allele-specific negative and positive controls. **C** Fold change in MFI of peptide-loaded monomers relative to allele-specific negative controls.

To identify CD8^+^ T cells that had responded to the vaccination, we first isolated single CD8^+^ T cells with an activated phenotype from PBMCs of five donors expressing at least one HLA allele with confirmed PfCSP peptide binding, before and after vaccination (Supp. Fig. 1D, 2A). TCR gene amplification and sequencing identified high frequencies of large persistent clones whose TCR genes showed similarity with those of TCRs with specificity against common viruses such as CMV, EBV, Influenza but also HBV, SARS-CoV, and YFV demonstrating that the activated CD8^+^ T cell repertoire under steady-state conditions is dominated by cells expressing TCR genes associated with anti-viral reactivity, which persist after vaccination (Supp. Fig. 2B, C). Expression cloning of 66 TCRs from cells that lacked these sequence features and were not detected before the vaccination failed to identify PfCSP-reactive clones demonstrating the relative scarcity of PfCSP-reactive CD8^+^ compared to anti-viral T cells with an activated phenotype (Supp Fig. 2D).

### Rare PfCSP-specific CD8^+^ T cells target an HLA class II restricted C-terminal epitope

To enrich for PfCSP-reactive CD8^+^ T cells, we performed *in vitro* stimulation experiments and assessed the frequency of activated CD8^+^ T cells (CD69^+^CD137^+^) analogous to the strategy successfully used for the enrichment of PfCSP-reactive CD4^+^ T cells (Supp Figure 1C). We initially stimulated PBMCs from two donors with peptides covering the complete PfCSP sequence, which led to strong activation and low levels of clonal expansion (Figure 5A, Supp Figure 1C). Stimulation of PBMCs from five additional donors with the separate peptide pools against the PfCSP subdomains induced C-CSP responses in four but N-CSP responses in only one donor (Figure 5A). In contrast to the N- or C-CSP specific clonal expansion observed for CD4^+^ T cell responses (Figure 1B-D), the majority of expanded CD8^+^ T cell clones (80%, F1, 100% 13/13, F3 62% 23/37) were detected across both peptide stimulation conditions indicative of bystander activation (Figure 5C). We therefore selected 30 expanded clones with V segments that were enriched uniquely after N- or C-CSP peptide pool stimulation without overlap between the two conditions for TCR expression cloning into CD8^+^ Jurkat76 T cells. Co-cultivation of the TCR transgenic cell lines with N-CSP and C-CSP peptide pool pulsed or unstimulated autologous B cells identified only one PfCSP-reactive clone with C-CSP specificity, whereas two clones responded under all conditions, most likely because they recognized abundant EBV peptides presented after the B cell immortalization (Figure 5D, Supp table 1).

**Figure 5:**
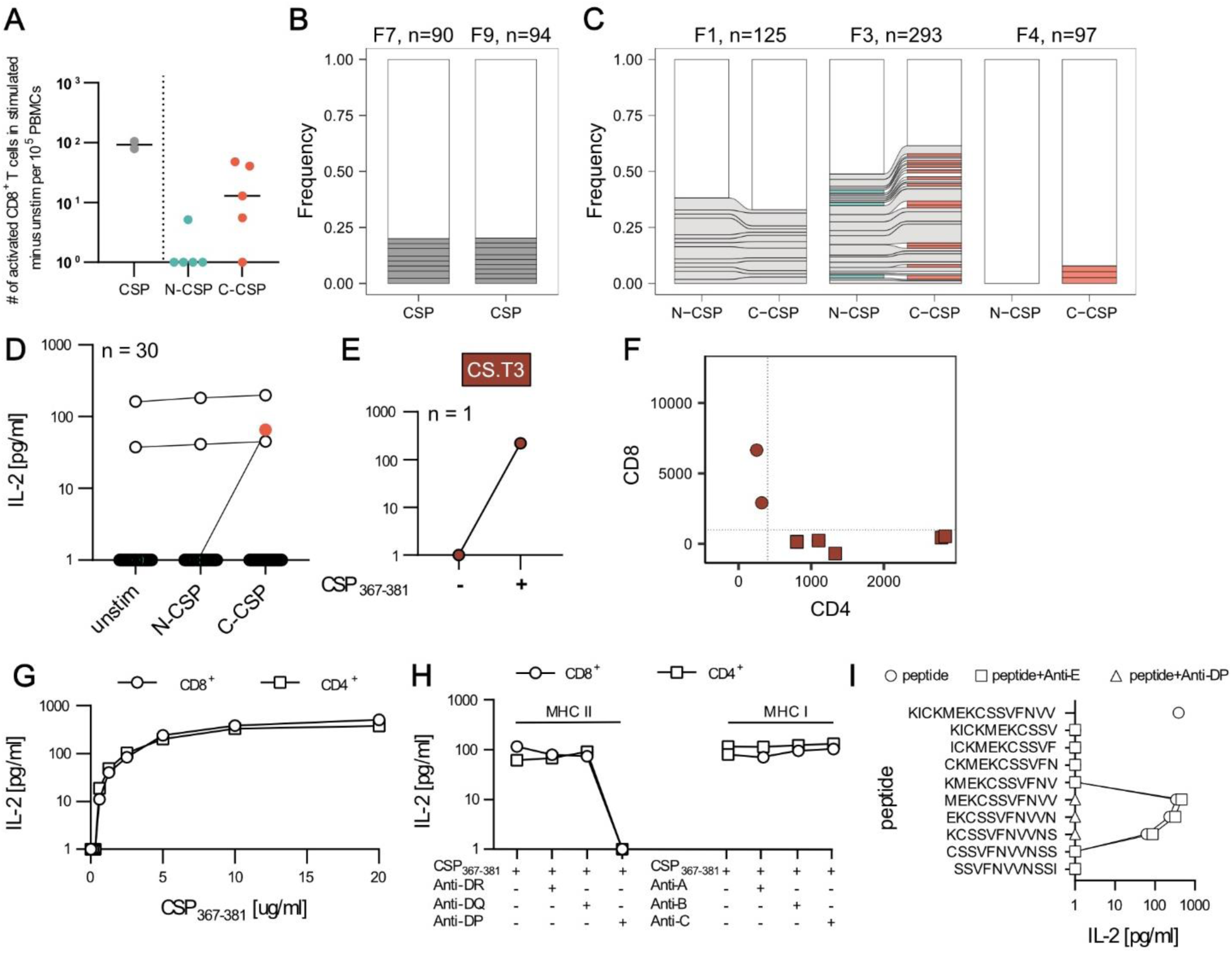
Rare PfCSP-specific CD8^+^ T cells target an HLA class II restricted C-terminal epitope co-receptor independently. **A** Number of activated (CD69^+^CD137^+^) CD8^+^ T cells from seven vaccinated donors (F7+F9, F1-5) after CSP, N-CSP, or C-CSP peptide pool mediated *in vitro* stimulation compared to unstimulated control cells from the same individuals. **B-C** Activated T cells were index single-cell sorted for paired *TRA* and *TRB* gene amplification and sequencing. Clonal composition of activated CD8^+^ T cells after CSP, N-CSP, or C-CSP peptide stimulation of samples from the indicated donors (F7, F9; F1, F3, F4). Expanded clones are shown in color and non-expanded clones are combined in white. Shared clones across separate expansion cultures are connected by ribbon. **D** Transgenic CD8^+^ Jurkat76 T cell lines were generated expressing TCRs from CD8^+^ T cells with an activated phenotype after stimulation of PBMCs with N- or C-CSP peptide pools. All monoclonal T cell lines were co-cultured with autologous B cells pulsed with N-CSP or C-CSP peptide pools or with non-peptide pulsed autologous B cells (unstimulated control). For each T cell line, the IL-2 concentrations in culture supernatants as measure for TCR-mediated activation were quantified by ELISA (individual dots). **E** C- and N-CSP peptide pool reactive TCR-transgenic Jurkat76 T cells were co-cultured with autologous B cells pulsed with individual C- or N-terminal peptides. IL-2 concentrations in culture supernatants were quantified by ELISA as measure for TCR-mediated activation. **F** CD4/CD8 phenotype of single-cell sorted T cells clonally related to the PfCSP-reactive T cell clone in the expansion culture. **G** IL-2 concentrations in supernatants of TCR-transgenic CD4^+^ or CD8^+^ Jurkat76 T cells co-cultured with CSP_367-381_ peptide-pulsed autologous B cells at the indicated peptide concentrations. **H** IL-2 concentrations in supernatants of TCR-transgenic CD4^+^ or CD8^+^ Jurkat76 T cells co-cultured with peptide-pulsed autologous B cells in the presence or absence of the indicated locus-specific HLA-blocking antibodies. **I** IL-2 concentrations in supernatants of TCR-transgenic CD8^+^ Jurkat76 T cells co-cultured with peptide-pulsed autologous B cells in the absence or presence of HLA-E or HLA-DP-specific HLA-blocking antibodies. **D,E,G,H** One out of two independent experiments is shown.

Stimulation with single peptide pulsed autologous B cells identified peptide 367-381 representing CS.T3 as cognate epitope of the C-CSP reactive CD8^+^ TCR (Figure 5E). Because our data showed that the CS.T3 epitope was also recognized by CD4^+^ T cells (Figure 2), we assessed the phenotype of the cell from which the TCR had been cloned. The CS.T3-reactive CD8^+^ T cell belonged to a larger clone of six members including four with a CD4^+^ phenotype, suggesting that this TCR binds the CS.T3 epitope co-receptor independently (Figure 5F). Indeed, when we expressed the TCR in CD4^+^ Jurkat76 T cells we observed the same concentration-dependent activation kinetic after CS.T3 peptide stimulation as with the CD8^+^ Jurkat76 T cells, demonstrating the co-receptor independence of this TCR (Figure 5G). Surprisingly, the use of anti-HLA blocking antibodies showed that the activation of both cell lines was blocked by the addition of an HLA-DP-specific antibody demonstrating that the TCR recognized the peptide in the context of class II even when expressed by CD8^+^ T cells (Figure 5H). Using an HLA-E specific blocking antibody we excluded the possibility that recognition of the CS.T3 peptide 367-381 was HLA-E dependent, although the peptide was predicted to be a strong HLE-E binder (Figure 5I). Through HLA-typing and peptide binding predictions we linked CS.T3 targeting to two non-shared HLA-DP alleles, HLA-DPB*10:01 and HLA-DPB*17:01, with high predicted binding to the core peptide EKCSSVFNVVN. The TCR of the CS.T3-reactive clone with mixed CD8^+^ and CD4^+^ phenotype was encoded by *TRBV7-2* and *TRAV36DV7* and thus showed no similarity with the CS.T3 specific TCRs of cells with only a CD4^+^ phenotype which were frequently encoded by *TRBV20-1* (Fig. 2) adding to the uniqueness of the co-receptor independent TCR.

In summary, although we were unable to identify class I restricted TCRs with reactivity against any of the predicted PfCSP peptides with strong binding to diverse HLA-A and HLA-B molecules, we identified rare CD8^+^ T cells with specificity for CS.T3 that recognized their target peptide co-receptor independently when presented on HLA-DP, just like their CD4^+^ clonal relatives.

Previous studies on T cell responses to PfCSP primarily focused on highly immunogenic but polymorphic CD4^+^ T cell epitopes, leaving subdominant epitopes with high sequence conversation largely uncharacterized (Moreno et al., 1993). Using human blood samples from FMP013/ALFQ vaccinees, we detected novel conserved epitopes in the PfCSP N- and C terminus which had not been identified after RTS,S or irradiated sporozoite vaccination (Lalvani et al., 1999; Wahl et al., 2022b). Although *in vitro* stimulation enabled the amplification and detection of T cell responses to less immunogenic epitopes, the high degree of non PfCSP-specific bystander activation and substantial overlap between CD4^+^ and CD8^+^ T cell epitopes highlights the need for functional validation at monoclonal TCR level or at least subset-specific T cell depletion (Oberhardt et al., 2021; Gondré-Lewis et al., 2023; Montes et al., 2005; Reiss et al., 2017).

The stronger cellular response and higher frequency of C-CSP compared to N-CSP peptide pool reactive T cells in all donors confirms the immunodominance of the C over the N terminus and junction region, similar to responses observed with irradiated sporozoite vaccination (Wahl et al., 2022b). Nonetheless, FMP013/ALFQ facilitated responses against novel N-terminal epitopes (CSP61, CSP81). Whether the PfCSP N terminus may represent an underappreciated target for T cell-based immunity requires confirmation. Our previous studies failed to identify N terminus reactive T cells in individuals vaccinated with high doses of irradiated sporozoites, potentially linked to the proteolytic cleavage of the PfCSP N terminus on the parasite surface or the fact that we did not enrich for N terminus reactivity by *in vitro* stimulation (Coppi et al., 2011; Singer et al., 2024).

While FMP013/ALFQ vaccination induced responses against all reported PfCSP CD4^+^ T cell epitopes, the CD8^+^ T cell response was restricted to CS.T3, contrasting the high number of CD8^+^ epitopes across all subdomains that have been reported in the past and expectations based on mouse studies (Heide et al., 2019). This discrepancy might be explained by the fact that the historic PfCSP CD8^+^ T cell epitope and HLA restrictions data originate from bulk cell stimulation and peptide binding predictions or *in vitro* peptide binding assays that were never confirmed at monoclonal TCR level (Sedegah et al., 2013; Heide et al., 2019). Alternatively, PfCSP, as delivered in the FMP013/ALFQ vaccine, may not effectively prime CD8^+^ T cells in humans compared to sporozoite immunization or natural *Pf* infection. Nonetheless, we identified a rare CS.T3-specific CD8^+^ T cell clone suggesting that this epitope is a relevant target of the cytotoxic anti-PfCSP response.

Our analysis revealed associations between TCR V-gene usage and epitope specificity, including the strong enrichment of TRBV30-1 in T1-specific TCRs and TRBV20-1 in CS.T3-specific TCRs. These findings align with previous reports of conserved TCR motifs linked to PfCSP-specific responses and together might provide an opportunity to identify T cell responses against defined epitopes by sequence analysis alone (Wahl et al., 2022b). Furthermore, our data illustrate the positive correlation between the diversity of HLAs that present a given peptide and the strength of the T cell response against this target. The fact that the highly immunogenic Th2R and T* epitopes lie in the most polymorphic regions of PfCSP, likely reflect the immune pressure that cellular responses exert on natural parasite populations. The conserved nature of the weaker immunogenic CSP61, CSP81, T1 and CS.T3 epitopes enhances their utility as vaccine targets for inducing strain-transcending immunity, addressing a critical limitation of RTS,S/AS01, which exhibits reduced efficacy against natural parasite populations with little sequence homology to the 3D7-derived vaccine C-terminal domain (Neafsey et al., 2015). Nevertheless, their stronger HLA restriction poses a challenge to the development of improved vaccine candidates that will be capable of inducing broad cellular anti-PfCSP immune responses.

The most promising seems to be CS.T3, which binds to diverse HLA alleles (Sinigaglia et al., 1988b), was target by CD4^+^ and CD8^+^ T cells, and is associated with protective T cell responses to natural infections (Reece et al., 2004). The high degree of sequence conservation likely reflects a yet unknown critical function of this PfCSP epitope for parasite development in the mosquito vector or human host. The low number of CS.T3 responders after RTS,S and irradiated sporozoite vaccination (Lalvani et al., 1999; Wahl et al., 2022b) and scarcity of the CS.T3-presenting HLA-DRB*10 allele in this study highlight the challenge of inducing CS.T3-specific CD4^+^ T cell responses by vaccination. The unique anti-CS.T3 T cell clone with CD4^+^ and CD8^+^ members that we identified recognized its peptide exclusively in the context of HLA-DP indicating a dominant role of CD4^+^ T cells in CS.T3 targeting. The co-receptor independent activation suggests high TCR affinity to compensate for the lack of co-receptor signaling in the context of CD8^+^ T cells, (Laugel et al., 2007), potentially enhancing cellular responses through the simultaneous activation of functionally diverse CD4^+^ and CD8^+^ T cell subsets (Davari et al., 2021). This unique TCR configuration highlights the potential for unconventional PfCSP-specific T cell responses that may bypass classical MHC restrictions.

Recent findings highlight the potential of CD4^+^ T cells to exhibit cytotoxic activity and produce high levels of IFN-γ (Moreno et al., 1991; Oliveira et al., 2008), challenging the traditional association of cytotoxic function with CD8^+^ T cells. While this study characterized the T cell response to FMP013/ALFQ vaccination, recently advancing alternative vaccine strategies including genetically-attenuated sporozoite vaccination and nucleic acid-based vaccines might lead to the induction of higher antigen-specific T cell frequencies through improved MHC presentation and T cell priming (Hoffman and Doolan, 2000; Wang et al., 1998; Lamers et al., 2024). It will be crucial to evaluate those T cell responses at monoclonal TCR level in order to gain additional information on gene sequence features associated with HLA and peptide binding and obtain training sets for algorithms that reliably predict T cell responses against PfCSP epitopes in malaria vaccine target populations. Although conserved PfCSP epitopes are restricted to less prevalent HLA alleles, combinations of epitopes that are presented on several low prevalent HLA alleles can reach population-wide coverage lowering the risk for T cell-mediated escape (Tsuji and Zavala, 2001).

This study has several limitations. First, the analysis was restricted to malaria-naive individuals in the US, and it remains unclear how prior exposure to *Pf* and HLA haplotype diversity in individuals living in malaria endemic or other infections might influence PfCSP-specific T cell responses. Second, while we identified several conserved epitopes and their HLA restrictions, functional validation of these responses *in vivo* and their correlation with protection remain to be established. Finally, the scarcity of PfCSP-specific CD8^+^ T cells highlights the need for further investigation into factors limiting their induction and the potential role of alternative PfCSP delivery platforms.

In summary, our study provides a detailed molecular characterization of PfCSP-specific T cell responses induced by FMP013/ALFQ vaccination providing a foundation for the rational design of vaccines that elicit robust, strain-transcending T cell responses. Future studies should focus on validating these findings in larger and more diverse cohorts and exploring strategies to enhance T cell priming for improved pre-erythrocytic immunity.

## Acknowledgments

We thank Christine Niesik, Yasmin Bergmann, Julia Gärtner, and Dorien Foster for experimental assistance, and Dr. Frank Momburg, Dr. Marten Meyer and Dr. Valerie Oberhardt for technical input and discussions. We are grateful to Dr. Theresa Kissel and Hendrik Feuerstein for critical reading of the manuscript and thank the vaccine trial participants and all members of the clinical trial unit at Walter Reed for their contribution and commitment to vaccine research. We thank Dr. Elke Bergmann-Leitner and the WRAIR Immunology Core Facility personnel for processing cells and serum samples from the FMP013 clinical trial. Clinical trial (NCT04268420) was conducted under a WRAIR IRB approved Protocol (2651) and the current study was approved by University of Heidelberg Ethics Committee (S-287/2021). The investigators have adhered to the policies for protection of human subjects as prescribed in AR 70–25. The material has been reviewed by the Walter Reed Army Institute of Research. There is no objection to its presentation and/or publication. The opinions or assertions contained herein are the private views of the author, and are not to be construed as official, or as reflecting true views of the Department of the Army, the Department of Defense.

---

**Supplementary figure 1:**
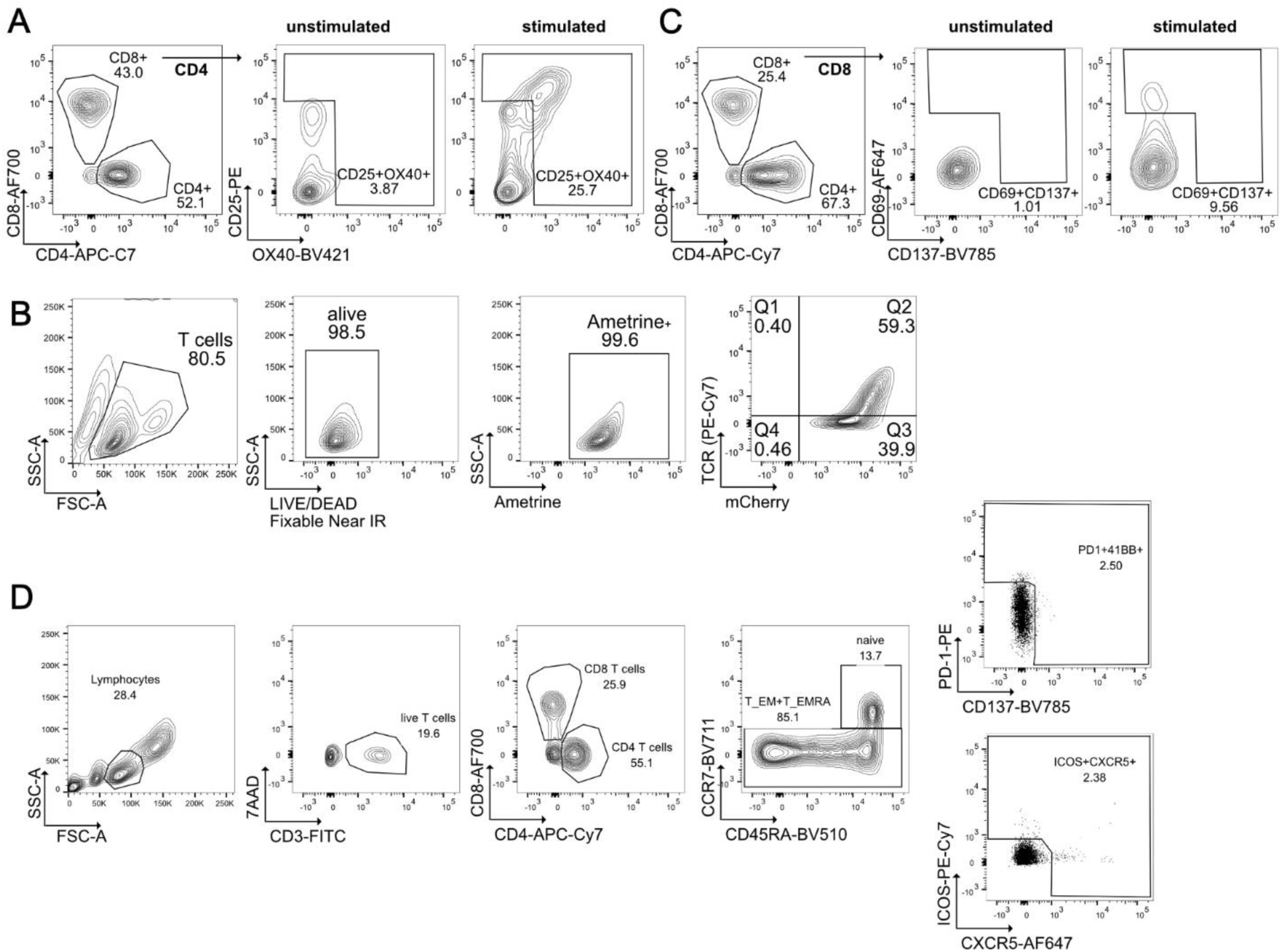
Gating strategy of activated CD4^+^/CD8^+^ T cells after *in vitro* expansion. **A, C** Gating strategy of activated CD4^+^ T cells (CD3^+^CD4^+^OX40^+^CD25^+^) (**A**) and CD8^+^ T cells (CD3^+^CD8^+^CD137^+^CD69^+^) (**C**) pre-gated on viable (7AAD-) lymphocytes (SSC/FSC) after *in vitro* stimulation with PfCSP peptides or in unstimulated control culture. **B** Gating strategy of TCR-transgenic Jurkat76 cell lines based on mCherry (mCherry^+^) and TCR (TCR^+^) expression quantified by flow cytometric analysis. **D** Gating strategy for flow cytometric analysis of PBMC samples from individuals receiving FMP013/ALFQ vaccinations.

**Supplementary figure 2:**
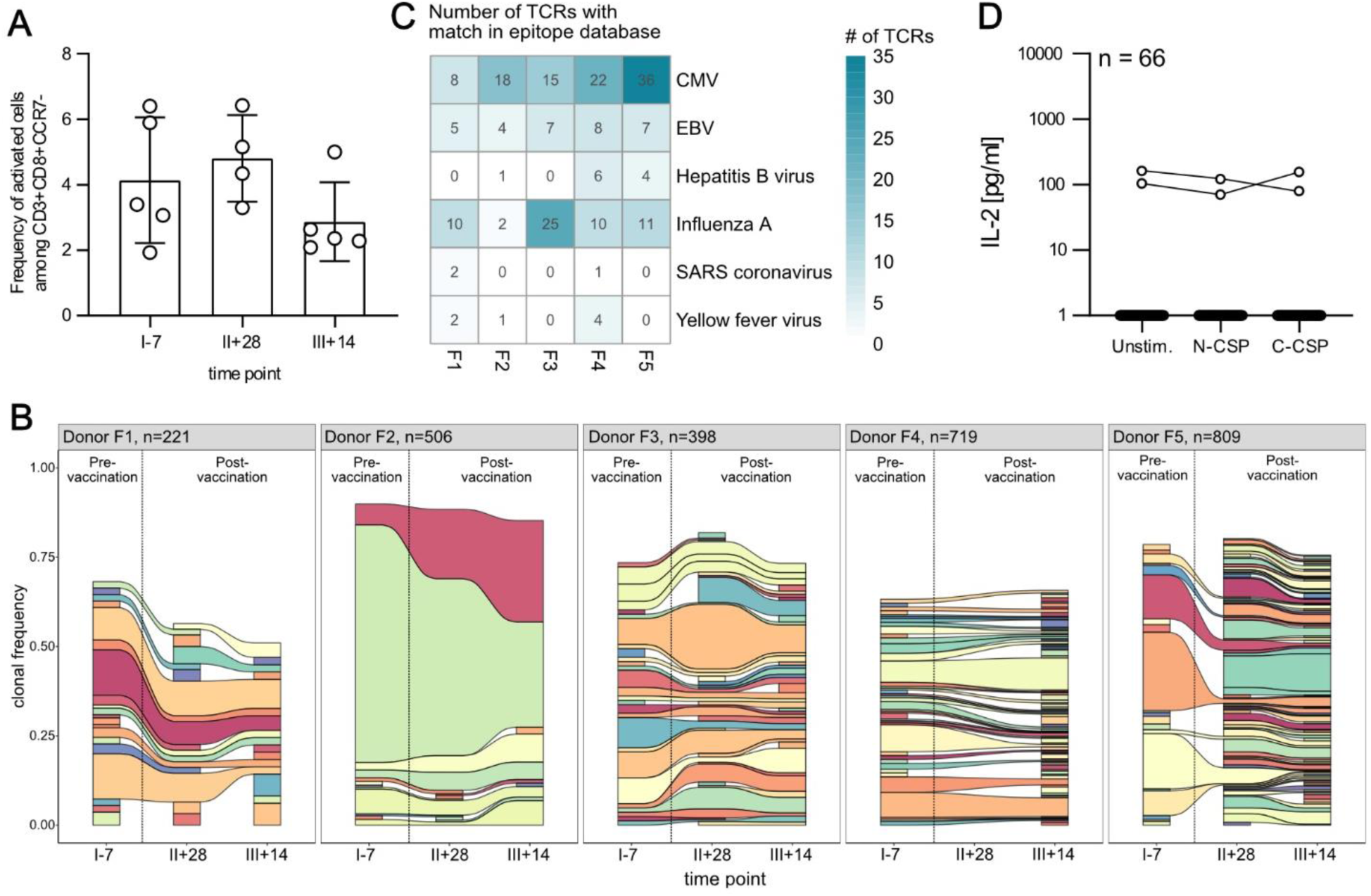
The activated CD8^+^ T cell repertoire is dominated by large clones with sequence similarity to T cell clones against common viruses. **A** Frequency of activated (PD1^+^CD137^+^ICOS^+^CXCR5^+^) T effector memory (T_EM_) or terminally differentiated T effector memory (T_EMRA_) cells as frequency of CD3^+^CD8^+^CCR7^−^ T cells for five donors (F1-5). Bar plots of individual timepoints before first (I-7), post second (II+28), and post third (III+14) vaccine dose show mean and standard deviation with individual dots representing individual donors. **B** Activated CD8^+^ T_EM+EMRA_ cells were indexed single-cell sorted, *TR* genes were amplified and sequenced for subsequent repertoire analysis. Clonal composition of the TCR gene repertoire in individual donors across multiple blood collection time points. Expanded clones are shown in color while unique TCR sequences are combined in the white compartment. Same color within each donor represents the same clone, while color sharing across different donors does not. **C** Number of TCRs with TCR beta chain match in IEDB identified by TCRmatch tool (Chronister et al., 2021). Identified TCRs are stratified by donor (F1-F5) and origin of the target epitope. CMV: Cytomegalovirus, EBV: Epstein-Barr virus, SARS: Severe acute respiratory syndrome **D** Transgenic CD8^+^ Jurkat76 T cell lines expressing TCRs from T cell clones without TCR sequence features of common virus-specific TCRs or presence before vaccination were generated. IL-2 concentrations in supernatants of 67 TCR-transgenic CD8^+^ Jurkat76 T cells co-cultured with autologous B cells pulsed with N-CSP or C-CSP peptide pools or with non-peptide-pulsed autologous B cells (unstimulated control).

**Supplementary table 1:** Sequence features, TCR expression, and reactivity of cloned and expressed TCRs.

| TCR ID | Donor | Time point | Population | TRBV | TRBJ | TRBC | TRAV | TRAJ | TCR exp. [%] | Reactivity |
| --- | --- | --- | --- | --- | --- | --- | --- | --- | --- | --- |
| T21F7x1604 | F7 | II+28 | CD4 | 7-2 | 2-7 | 2 | 10-1 | 18 | 32.7 | Non-CSP-reactive |
| T21F9x2086 | F9 | II+28 | CD4 | 6-1 | 2-7 | 2 | 14DV4 | 13 | 63 | CS.T3 |
| T21F7x1777 | F7 | II+28 | CD4 | 7-2 | 2-1 | 2 | 10-1 | 18 | 56.4 | Non-CSP-reactive |
| T21F7x1643 | F7 | II+28 | CD4 | 7-2 | 2-7 | 2 | 10-1 | 18 | 66.8 | Non-CSP-reactive |
| T21F4x3646 | F4 | II+28 | CD4 | 20-1 | 2-5 | 2 | 17-1 | 50 | 41.5 | Non-CSP-reactive |
| T21F3x2835 | F3 | II+28 | CD4 | 12-3 | 1-5 | 1 | 17-1 | 11 | 71.6 | Th2R |
| T21F4x3818 | F4 | II+28 | CD4 | 19-1 | 2-7 | 2 | 13-1 | 47 | 27.8 | Non-CSP-reactive |
| T21F3x2797 | F3 | II+28 | CD4 | 5-1 | 2-6 | 2 | 12-1 | 20 | 42.8 | CSP61 |
| T21F3x2855 | F3 | II+28 | CD4 | 5-1 | 2-6 | 2 | 12-1 | 20 | 30.8 | CSP61 |
| T21F3x2742 | F3 | II+28 | CD4 | 5-1 | 2-6 | 2 | 12-1 | 20 | 69.5 | CSP61 |
| T21F3x2893 | F3 | II+28 | CD4 | 3-1 | 2-2 | 2 | 19-1 | 26 | 45.9 | CSP61 |
| T21F4x3685 | F4 | II+28 | CD4 | 20-1 | 2-2 | 2 | 9-2 | 49 | 78.9 | CS.T3 |
| T21F4x3722 | F4 | II+28 | CD4 | 27-1 | 1-4 | 1 | 1-2 | 33 | 69.7 | CSP81 |
| T21F7x1698 | F7 | II+28 | CD4 | 30-1 | 2-2 | 2 | 29DV5 | 18 | 64.3 | T* |
| T21F4x3701 | F4 | II+28 | CD4 | 20-1 | 2-5 | 2 | 9-2 | 57 | 68.4 | CS.T3 |
| T21F3x3023 | F3 | II+28 | CD4 | 14-1 | 1-2 | 1 | 19-1 | 43 | 46.9 | Non-CSP-reactive |
| T21F3x2714 | F3 | II+28 | CD4 | 30-1 | 1-5 | 1 | 24-1 | 28 | 75.3 | T1 |
| T21F3x2705 | F3 | II+28 | CD4 | 30-1 | 1-3 | 1 | 24-1 | 32 | 68.9 | T1 |
| T21F3x2851 | F3 | II+28 | CD4 | 30-1 | 1-6 | 1 | 3-1 | 36 | 81 | T1 |
| T21F3x2777 | F3 | II+28 | CD4 | 30-1 | 1-2 | 1 | 36DV7 | 54 | 66.1 | T1 |
| T21F3x2838 | F3 | II+28 | CD4 | 12-3 | 1-5 | 1 | 17-1 | 11 | 66.7 | Th2R |
| T21F3x2968 | F3 | II+28 | CD4 | 12-4 | 1-2 | 1 | 17-1 | 33 | 28.8 | Non-CSP-reactive |
| T21F3x2862 | F3 | II+28 | CD4 | 6-4 | 1-2 | 1 | 17-1 | 11 | 67.6 | Th2R |
| T21F4x3702 | F4 | II+28 | CD4 | 20-1 | 2-7 | 2 | 9-2 | 58 | 65.4 | CS.T3 |
| T21F3x2939 | F3 | II+28 | CD4 | 12-3 | 1-3 | 1 | 26-1 | 48 | 76.1 | Non-CSP-reactive |
| T21F3x3026 | F3 | II+28 | CD4 | 7-2 | 2-3 | 2 | 26-1 | 23 | 53.3 | Non-CSP-reactive |
| T21F4x3706 | F4 | II+28 | CD4 | 20-1 | 2-1 | 2 | 9-2 | 10 | 86 | CS.T3 |
| T21F3x2949 | F3 | II+28 | CD4 | 12-3 | 1-2 | 1 | 23DV6 | 23 | 43.3 | Non-CSP-reactive |
| T21F3x2863 | F3 | II+28 | CD4 | 7-2 | 1-5 | 1 | 17-1 | 11 | 56 | Th2R |
| T21F3x2903 | F3 | II+28 | CD4 | 7-2 | 2-1 | 2 | 17-1 | 11 | 49.5 | Th2R |
| T21F3x2996 | F3 | II+28 | CD4 | 5-1 | 1-5 | 1 | 29DV5 | 40 | 71.8 | Non-CSP-reactive |
| T21F3x2911 | F3 | II+28 | CD4 | 18-1 | 1-2 | 1 | 13-1 | 53 | 72.2 | Th2R |
| T21F3x2865 | F3 | II+28 | CD4 | 7-2 | 2-3 | 2 | 35-1 | 17 | 38.8 | Non-CSP-reactive |
| T21F3x3008 | F3 | II+28 | CD4 | 27-1 | 1-6 | 1 | 26-2 | 40 | 60.1 | Non-CSP-reactive |
| T21F4x3724 | F4 | II+28 | CD4 | 20-1 | 2-2 | 2 | 35-1 | 49 | 56.4 | Non-CSP-reactive |
| T21F7x1635 | F7 | II+28 | CD4 | 5-8 | 1-1 | 1 | 29DV5 | 45 | 58.5 | Non-CSP-reactive |
| T21F3x2915 | F3 | II+28 | CD4 | 7-2 | 1-5 | 1 | 17-1 | 11 | 41.5 | Th2R |
| T21F9x2073 | F9 | II+28 | CD4 | 6-1 | 1-5 | 1 | 29DV5 | 53 | 64 | Non-CSP-reactive |
| T21F9x2018 | F9 | II+28 | CD4 | 12-4 | 2-7 | 2 | 39-1 | 48 | 61.1 | Non-CSP-reactive |
| T21F4x3758 | F4 | II+28 | CD4 | 28-1 | 2-1 | 2 | 41-1 | 57 | 24 | Non-CSP-reactive |
| T21F4x3762 | F4 | II+28 | CD4 | 20-1 | 2-5 | 2 | 9-2 | 35 | 68.4 | CS.T3 |
| T21F4x3653 | F4 | II+28 | CD4 | 28-1 | 2-1 | 2 | 35-1 | 28 | 60.3 | Non-CSP-reactive |
| T21F3x2917 | F3 | II+28 | CD4 | 7-7 | 1-1 | 1 | 8-4 | 15 | 34.6 | Th2R |
| T21F3x2907 | F3 | II+28 | CD4 | 5-1 | 2-1 | 2 | 38-2-DV8 | 52 | 75 | Non-CSP-reactive |
| T21F3x2931 | F3 | II+28 | CD4 | 3-1 | 1-3 | 1 | 17-1 | 34 | 59.2 | Th2R |
| T21F3x2795 | F3 | II+28 | CD4 | 3-1 | 2-4 | 2 | 8-3 | 45 | 12 | Non-CSP-reactive |
| T21F3x2945 | F3 | II+28 | CD4 | 12-3 | 1-5 | 1 | 17-1 | 11 | 63.9 | Th2R |
| T21F3x3020 | F3 | II+28 | CD4 | 12-3 | 1-1 | 1 | 8-4 | 50 | 76 | Non-CSP-reactive |
| T21F4x3712 | F4 | II+28 | CD4 | 5-4 | 2-7 | 2 | 8-4 | 49 | 63.2 | Non-CSP-reactive |
| T21F3x3018 | F3 | II+28 | CD4 | 14-1 | 1-2 | 1 | 9-2 | 20 | 70.3 | Non-CSP-reactive |
| T21F3x2960 | F3 | II+28 | CD4 | 12-3 | 1-5 | 1 | 17-1 | 11 | 71.5 | Th2R |
| T21F4x3791 | F4 | II+28 | CD4 | 20-1 | 2-1 | 2 | 9-2 | 32 | 79.8 | Non-CSP-reactive |
| T21F3x2975 | F3 | II+28 | CD4 | 5-1 | 2-5 | 2 | 9-2 | 52 | 71.6 | Th2R |
| T21F4x3795 | F4 | II+28 | CD4 | 20-1 | 1-4 | 1 | 13-1 | 29 | 76.2 | EBV |
| T21F3x3019 | F3 | II+28 | CD4 | 5-1 | 2-5 | 2 | 9-2 | 52 | 69 | Th2R |
| T21F4x3804 | F4 | II+28 | CD4 | 20-1 | 2-1 | 2 | 12-3 | 45 | 68 | Non-CSP-reactive |
| T21F4x3682 | F4 | II+28 | CD4 | 20-1 | 2-4 | 2 | 13-1 | 53 | 61.8 | Th2R |
| T21F3x3049 | F3 | II+28 | CD4 | 30-1 | 1-1 | 1 | 13-2 | 31 | 70.4 | Non-CSP-reactive |
| T21F3x2731 | F3 | II+28 | CD4 | 30-1 | 1-5 | 1 | 23DV6 | 36 | 69.3 | T1 |
| T21F3x2717 | F3 | II+28 | CD4 | 30-1 | 2-7 | 2 | 36DV7 | 54 | 77.1 | T1 |
| T21F3x2989 | F3 | II+28 | CD4 | 5-1 | 2-5 | 2 | 9-2 | 52 | 74.9 | Th2R |
| T21F3x2898 | F3 | II+28 | CD4 | 5-1 | 2-6 | 2 | 9-2 | 49 | 82.5 | Non-CSP-reactive |
| T21F3x2807 | F3 | II+28 | CD4 | 30-1 | 2-3 | 2 | 26-1 | 27 | 74.2 | Non-CSP-reactive |
| T21F4x3806 | F4 | II+28 | CD4 | 20-1 | 2-2 | 2 | 8-3 | 15 | 31.4 | Non-CSP-reactive |
| T21F4x3825 | F4 | II+28 | CD4 | 20-1 | 2-5 | 2 | 9-2 | 10 | 68.9 | CS.T3 |
| T21F3x2801 | F3 | II+28 | CD4 | 20-1 | 2-7 | 2 | 13-1 | 15 | 61.7 | T* |
| T19F1x1257 | F1 | II+28 | T<sub>EM</sub>+T<sub>EMRA</sub> | 27 | 2-1 | 2 | 2 | 30 | 16 | Non-CSP-reactive |
| T19F1x2356 | F1 | III+14 | T<sub>EM</sub>+T<sub>EMRA</sub> | 12-3 | 1-1 | 1 | 29/DV5 | 43 | 58 | Non-CSP-reactive |
| T19F1x2391 | F1 | III+14 | T<sub>EM</sub>+T<sub>EMRA</sub> | 11-2 | 2-1 | 2 | 13-2 | 47 | 24 | Non-CSP-reactive |
| T19F2x1302 | F2 | II+28 | T<sub>EM</sub>+T<sub>EMRA</sub> | 27 | 2-7 | 2 | 12-3 | 13 | 71 | EBV reactive |
| T19F3x874 | F3 | III+14 | T<sub>EM</sub>+T<sub>EMRA</sub> | 7-9 | 1-6 | 1 | 41 | 57 | 41 | Non-CSP-reactive |
| T19F3x906 | F3 | III+14 | T<sub>EM</sub>+T<sub>EMRA</sub> | 27 | 2-5 | 2 | 29/DV5 | 47 | 45 | Non-CSP-reactive |
| T19F4x3235 | F4 | III+14 | T<sub>EM</sub>+T<sub>EMRA</sub> | 4-3 | 2-2 | 2 | 12-2 | 49 | 29 | Non-CSP-reactive |
| T19F4x3308 | F4 | III+14 | T<sub>EM</sub>+T<sub>EMRA</sub> | 12-3 | 1-2 | 1 | 12-2 | 49 | 69 | Non-CSP-reactive |
| T19F4x3320 | F4 | III+14 | T<sub>EM</sub>+T<sub>EMRA</sub> | 12-3 | 1-5 | 1 | 12-3 | 49 | 47 | Non-CSP-reactive |
| T19F5x1657 | F5 | III+14 | T<sub>EM</sub>+T<sub>EMRA</sub> | 18 | 1-1 | 1 | 41 | 48 | 61 | Non-CSP-reactive |
| T19F5x1793 | F5 | III+14 | Tet+ | 28 | 1-6 | 1 | 26-2 | 48 | 83 | Non-CSP-reactive |
| T19F5x1850 | F5 | III+14 | Tet+ | 28 | 2-3 | 2 | 12-2 | 45 | 59 | Non-CSP-reactive |
| T19F5x2130 | F5 | III+14 | T<sub>EM</sub>+T<sub>EMRA</sub> | 28-1 | 2-1 | 2 | 12-1 | 49 | 42 | Non-CSP-reactive |
| T19F5x2143 | F5 | III+14 | T<sub>EM</sub>+T<sub>EMRA</sub> | 7-6 | 1-5 | 1 | 26-1 | 49 | 60 | Non-CSP-reactive |
| T19F5x2201 | F5 | III+14 | Tet+ | 9-1 | 2-1 | 2 | 13-1 | 15 | 63 | Non-CSP-reactive |
| T19F5x2242 | F5 | III+14 | Tet+ | 7-9 | 1-6 | 1 | 8-2 | 12 | 25 | Non-CSP-reactive |
| T19F5x2245 | F5 | III+14 | Tet+ | 28-1 | 2-3 | 2 | 12-1 | 49 | 63 | Non-CSP-reactive |
| T19F5x2251 | F5 | III+14 | Tet+ | 27 | 1-2 | 1 | 19 | 44 | 52 | Non-CSP-reactive |
| T19F5x4251 | F5 | II+28 | T<sub>EM</sub>+T<sub>EMRA</sub> | 20-1 | 2-7 | 2 | 27 | 24 | 50 | Non-CSP-reactive |
| T19F5x4261 | F5 | II+28 | T<sub>EM</sub>+T<sub>EMRA</sub> | 28 | 2-3 | 2 | 12-2 | 49 | 54 | Non-CSP-reactive |
| T19F5x4313 | F5 | II+28 | T<sub>EM</sub>+T<sub>EMRA</sub> | 20-1 | 2-5 | 2 | 30 | 12 | 22 | Non-CSP-reactive |
| T19F5x4355 | F5 | II+28 | T<sub>EM</sub>+T<sub>EMRA</sub> | 25-1 | 1-3 | 1 | 13-2 | 16 | 34 | Non-CSP-reactive |
| T19F5x4356 | F5 | II+28 | T<sub>EM</sub>+T<sub>EMRA</sub> | 28 | 1-6 | 1 | 12-2 | 28 | 67 | Non-CSP-reactive |
| T19F5x4361 | F5 | II+28 | T<sub>EM</sub>+T<sub>EMRA</sub> | 12-4 | 2-3 | 2 | 12-2 | 49 | 64 | Non-CSP-reactive |
| T19F5x4395 | F5 | II+28 | T<sub>EM</sub>+T<sub>EMRA</sub> | 5-8 | 2-7 | 2 | 14/DV4 | 8 | 69 | Non-CSP-reactive |
| T19F5x4452 | F5 | II+28 | Tet+ | 5-6 | 2-3 | 2 | 13-2 | 48 | 48 | Non-CSP-reactive |
| T19F5x4469 | F5 | II+28 | Tet+ | 27 | 1-4 | 1 | 21 | 26 | 69 | Non-CSP-reactive |
| T19F5x4473 | F5 | II+28 | Tet+ | 7-6 | 1-4 | 1 | 26-2 | 43 | 66 | Non-CSP-reactive |
| T19F5x4501 | F5 | II+28 | Tet+ | 9 | 1-5 | 1 | 19 | 30 | 41 | Non-CSP-reactive |
| T19F5x4523 | F5 | II+28 | Tet+ | 12-4 | 2-7 | 2 | 1-1 | 41 | 25 | Non-CSP-reactive |
| T19F5x4529 | F5 | II+28 | Tet+ | 27 | 2-1 | 2 | 21 | 26 | 63 | Non-CSP-reactive |
| T19F5x4850 | F5 | II+28 | Tet+ | 19-1 | 2-1 | 2 | 12-2 | 42 | 28 | Non-CSP-reactive |
| T19F5x4886 | F5 | II+28 | Tet+ | 7-8 | 2-4 | 2 | 21-1 | 49 | 47 | Non-CSP-reactive |
| T19F5x4909 | F5 | II+28 | Tet+ | 20-1 | 2-7 | 2 | 8-6 | 52 | 62 | Non-CSP-reactive |
| T19F5x4912 | F5 | II+28 | Tet+ | 19-1 | 1-5 | 1 | 12-2 | 30 | 57 | Non-CSP-reactive |
| T20F2x2439 | F2 | III+14 | T<sub>EM</sub>+T<sub>EMRA</sub> | 5-8 | 2-5 | 2 | 13-2 | 45 | 52 | Non-CSP-reactive |
| T20F2x2549 | F2 | III+14 | Tet+ | 10-2 | 2-7 | 2 | 12-1 | 37 | 53 | Non-CSP-reactive |
| T20F3x2177 | F3 | II+28 | T<sub>EM</sub>+T<sub>EMRA</sub> | 2-1 | 2-7 | 2 | 13-1 | 24 | 62 | Non-CSP-reactive |
| T20F4x2630 | F4 | III+14 | T<sub>EM</sub>+T<sub>EMRA</sub> | 3-1 | 1-5 | 1 | 12-2 | 23 | 51 | Non-CSP-reactive |
| T20F4x2658 | F4 | III+14 | T<sub>EM</sub>+T<sub>EMRA</sub> | 12-3 | 2-7 | 2 | 26-1 | 20 | 68 | Non-CSP-reactive |
| T20F4x3312 | F4 | II+28 | CD8 | 7-9 | 1-2 | 1 | 29/DV5 | 52 | 30 | EBV reactive |
| T20F5x2724 | F5 | II+28 | Tet+ | 18-1 | 2-5 | 2 | 16-1 | 39 | 31 | Non-CSP-reactive |
| T20F5x2765 | F5 | II+28 | Tet+ | 6-2 | 2-7 | 2 | 10-1 | 34 | 35 | Non-CSP-reactive |
| T20F5x2820 | F5 | II+28 | Tet+ | 7-2 | 2-2 | 2 | 1-2 | 31 | 60 | Non-CSP-reactive |
| T20F5x2874 | F5 | II+28 | Tet+ | 5-6 | 2-3 | 2 | 4-1 | 5 | 67 | Non-CSP-reactive |
| T20F5x2913 | F5 | II+28 | T<sub>EM</sub>+T<sub>EMRA</sub> | 11-3 | 2-1 | 2 | 12-2 | 20 | 36 | Non-CSP-reactive |
| T20F5x2984 | F5 | II+28 | T<sub>EM</sub>+T<sub>EMRA</sub> | 3-1 | 2-1 | 2 | 8-4 | 54 | 55 | Non-CSP-reactive |
| T20F5x3019 | F5 | II+28 | T<sub>EM</sub>+T<sub>EMRA</sub> | 20-1 | 2-5 | 2 | 8-2 | 12 | 56 | Non-CSP-reactive |
| T20F5x3039 | F5 | II+28 | T<sub>EM</sub>+T<sub>EMRA</sub> | 15-1 | 2-1 | 2 | 22-1 | 35 | 57 | Non-CSP-reactive |
| T20F5x3873 | F5 | III+14 | T<sub>EM</sub>+T<sub>EMRA</sub> | 10-3 | 2-5 | 2 | 8-3 | 13 | 53 | Non-CSP-reactive |
| T20F5x3992 | F5 | III+14 | T<sub>EM</sub>+T<sub>EMRA</sub> | 6-5 | 1-2 | 1 | 4-1 | 41 | 73 | Non-CSP-reactive |
| T20F5x4028 | F5 | III+14 | T<sub>EM</sub>+T<sub>EMRA</sub> | 7-9 | 2-1 | 2 | 19-1 | 12 | 20 | Non-CSP-reactive |
| T20F5x4029 | F5 | III+14 | T<sub>EM</sub>+T<sub>EMRA</sub> | 19-1 | 2-7 | 2 | 19-1 | 42 | 36 | Non-CSP-reactive |
| T20F5x4035 | F5 | III+14 | T<sub>EM</sub>+T<sub>EMRA</sub> | 6-2 | 1-5 | 1 | 41-1 | 45 | 54 | Non-CSP-reactive |
| T20F5x4038 | F5 | III+14 | T<sub>EM</sub>+T<sub>EMRA</sub> | 3-1 | 1-1 | 1 | 38-2-DV8 | 40 | 63 | Non-CSP-reactive |
| T20F5x4097 | F5 | III+14 | Tet+ | 19-1 | 2-1 | 2 | 8-2 | 30 | 48 | Non-CSP-reactive |
| T20F5x4140 | F5 | III+14 | Tet+ | 18-1 | 2-7 | 2 | 12-1 | 44 | 37 | Non-CSP-reactive |
| T20F5x4162 | F5 | III+14 | Tet+ | 20-1 | 1-5 | 1 | 12-2 | 24 | 79 | Non-CSP-reactive |
| T20F5x4208 | F5 | III+14 | Tet+ | 6-2 | 1-1 | 1 | 12-2 | 12 | 55 | Non-CSP-reactive |
| T20F5x4215 | F5 | III+14 | Tet+ | 19-1 | 1-2 | 1 | 38-2-DV8 | 27 | 54 | Non-CSP-reactive |
| T19F4x2853 | F4 | III+14 | T<sub>EM</sub>+T<sub>EMRA</sub> | 12-3 | 1-2 | 1 | 12-1 | 31 | 70.8 | Non-CSP-reactive |
| T19F4x2853 | F4 | III+14 | T<sub>EM</sub>+T<sub>EMRA</sub> | 12-3 | 1-2 | 1 | 12-1 | 31 | 44.3 | Non-CSP-reactive |
| T19F5x4436 | F5 | II+28 | T<sub>EM</sub>+T<sub>EMRA</sub> | 28-1 | 2-1 | 2 | 1-2 | 27 | 13.7 | Non-CSP-reactive |
| T19F4x3016 | F4 | III+14 | T<sub>EM</sub>+T<sub>EMRA</sub> | 12-3 | 1-1 | 1 | 12-2 | 31 | 23.5 | Non-CSP-reactive |
| T19F3x915 | F3 | III+14 | Tet+ | 20-1 | 2-7 | 2 | 24-1 | 34 | 50.9 | Non-CSP-reactive |
| T19F5x4265 | F5 | II+28 | T<sub>EM</sub>+T<sub>EMRA</sub> | 27-1 | 2-2 | 2 | 27-1 | 50 | 36.6 | Non-CSP-reactive |
| T19F3x856 | F3 | III+14 | T<sub>EM</sub>+T<sub>EMRA</sub> | 12-4 | 1-2 | 1 | 35-1 | 50 | 38.5 | Non-CSP-reactive |
| T21F1x615 | F1 | II+28 | CD8 | 15-1 | 1-1 | 1 | 12-2 | 30 | 24.3 | Non-CSP-reactive |
| T21F3x649 | F3 | II+28 | CD8 | 12-4 | 1-1 | 1 | 29DV5 | 30 | 14.1 | Non-CSP-reactive |
| T21F3x661 | F3 | II+28 | CD8 | 5-1 | 2-7 | 2 | 25-1 | 32 | 56.2 | Non-CSP-reactive |
| T21F3x857 | F3 | II+28 | CD8 | 3-1 | 1-3 | 1 | 8-6 | 23 | 61.3 | Non-CSP-reactive |
| T21F3x863 | F3 | II+28 | CD8 | 20-1 | 2-7 | 2 | 24-1 | 34 | 78.6 | Non-CSP-reactive |
| T21F3x883 | F3 | II+28 | CD8 | 12-4 | 1-2 | 1 | 35-1 | 50 | 52.4 | Non-CSP-reactive |
| T21F3x1001 | F3 | II+28 | CD8 | 12-4 | 2-7 | 2 | 27-1 | 40 | 70.5 | Non-CSP-reactive |
| T21F3x1010 | F3 | II+28 | CD8 | 27-1 | 2-1 | 2 | 6-1 | 29 | 17.2 | Non-CSP-reactive |
| T21F3x1025 | F3 | II+28 | CD8 | 29-1 | 2-7 | 2 | 35-1 | 20 | 63 | Non-CSP-reactive |
| T21F3x1031 | F3 | II+28 | CD8 | 5-1 | 1-2 | 1 | 29DV5 | 40 | 29.5 | Non-CSP-reactive |
| T21F3x1037 | F3 | II+28 | CD8 | 12-4 | 2-4 | 2 | 22-1 | 29 | 55 | Non-CSP-reactive |
| T21F3x1053 | F3 | II+28 | CD8 | 5-5 | 1-3 | 1 | 22-1 | 48 | 77.6 | Non-CSP-reactive |
| T21F3x1101 | F3 | II+28 | CD8 | 5-1 | 1-2 | 1 | 17-1 | 53 | 61.4 | Non-CSP-reactive |
| T21F3x1113 | F3 | II+28 | CD8 | 5-1 | 1-1 | 1 | 19-1 | 57 | 59.2 | Non-CSP-reactive |
| T21F7x1757 | F7 | II+28 | CD8 | 30-1 | 2-2 | 2 | 12-2 | 15 | 57.3 | Non-CSP-reactive |
| T21F7x1811 | F7 | II+28 | CD8 | 19-1 | 2-1 | 2 | 12-1 | 8 | 30.9 | Non-CSP-reactive |
| T21F9x1889 | F9 | II+28 | CD8 | 7-2 | 1-2 | 1 | 36DV7 | 53 | 47 | CS.T3 |
| T21F7x1907 | F7 | II+28 | CD8 | 30-1 | 2-2 | 2 | 12-2 | 15 | 62.9 | Non-CSP-reactive |
| T21F9x2160 | F9 | II+28 | CD8 | 7-2 | 1-4 | 1 | 1-2 | 20 | 66.6 | Non-CSP-reactive |
| T21F7x2213 | F7 | II+28 | CD8 | 7-9 | 2-1 | 2 | 12-2 | 24 | 34.2 | Non-CSP-reactive |
| T21F7x2225 | F7 | II+28 | CD8 | 5-8 | 2-3 | 2 | 29DV5 | 45 | 51.2 | Non-CSP-reactive |
| T21F4x2344 | F4 | II+28 | CD8 | 5-1 | 2-7 | 2 | 1-2 | 33 | 69.4 | Non-CSP-reactive |
| T21F4x2345 | F4 | II+28 | CD8 | 20-1 | 2-1 | 2 | 26-1 | 9 | 64.6 | Non-CSP-reactive |
| T21F4x2376 | F4 | II+28 | CD8 | 4-1 | 1-2 | 1 | 8-3 | 35 | 13.1 | Non-CSP-reactive |
| T21F4x2396 | F4 | II+28 | CD8 | 19-1 | 2-7 | 2 | 22-1 | 24 | 58.3 | Non-CSP-reactive |
| T21F4x2402 | F4 | II+28 | CD8 | 20-1 | 2-2 | 2 | 39-1 | 16 | 49.9 | Non-CSP-reactive |
| T21F4x2507 | F4 | II+28 | CD8 | 12-3 | 2-2 | 2 | 1-2 | 33 | 68.7 | Non-CSP-reactive |
| T21F4x2512 | F4 | II+28 | CD8 | 20-1 | 2-7 | 2 | 13-1 | 13 | 46.8 | Non-CSP-reactive |
| T21F4x2528 | F4 | II+28 | CD8 | 5-1 | 2-1 | 2 | 17-1 | 57 | 51.7 | Non-CSP-reactive |
| T21F4x2494 | F4 | II+28 | CD8 | 27-1 | 1-2 | 1 | 1-2 | 6 | 61 | Non-CSP-reactive |

## Methods

### Human FMP013 vaccine trial

PBMC samples from participants of the Falciparum Malaria Protein 013 (FMP013) vaccination trial (ClinicalTrials.gov number NCT04268420) isolated 7 days before the first immunization, 28 days after the second, and 14 days after the third immunization were analyzed upon approval by the ethics committee of the medical faculty of the University Heidelberg (number: S-287/2021). Study participants were healthy volunteers with no history of malaria or HIV, who had received three doses of either 20 µg (F6-F10) or 40 µg (F1-F5) FMP013 in ALFQ adjuvant on days 1, 29, and 57.

### *In vitro* T cell expansion

PBMCs were thawed and resuspended in 5 ml RPMI containing 10 % fetal calf serum (FCS, US origin), 2 mM Glutamine, 1.2 % Penicillin/Streptomycin and 1.5 % 1M HEPES (expansion medium). Cells were divided into three aliquots and 1×10^6^ cells were incubated with the C-CSP or N-CSP peptide pools (15mer peptides, 11aa overlap N-CSP: aa 20-281; C-CSP: aa271-385) or an equivalent amount of DMSO (unstimulated control), respectively. After incubation for 1 h at 37 °C, cells were washed with 5 ml RPMI, combined with 2×10^6^ unstimulated cells and seeded into a 48-well cell culture plate at densities of 1.5×10^6^ cells per well. To provide co-stimulatory signals, 20 U/ml recombinant IL-2 (Stemcell) and 0.5 µg/ml CD28 antibody (BD Biosciences) were added to the culture. On days 4 and 7, the expansion medium was exchanged and cells were transferred to a 24-well cell culture plate according to the cell density. On day 9, cells were starved for 24 h in medium without IL-2 supplementation to minimize unspecific activation.

### Flow cytometry and single-cell sorting

Ten-day stimulated and expanded cells were incubated for 30 min at 4°C in the dark with 100 µl of antibody staining cocktail, containing the following antibodies diluted in fluorescence-activated cell sorting (FACS) buffer (4% fetal bovine serum (FBS) in PBS): CD3-FITC (Biolegend, catalog no. 317306), CD4-APC-Cy7 (Biolegend, catalog no. 357416), CD8a-Alexa Fluor 700 (BioLegend, catalog no. 344724), CD137-BV785 (BD Biosciences, cat. no. 744397), CD69-Alexa Fluor647 (Biolegend, catalog no. 310918), OX40-BV421 (Biolegend, cat. no. 350013), CD25-PE (Biolegend, catalog no. 302606). Freshly thawed and washed PBMCs were incubated for 30 min at 4°C in the dark with 100 µl of antibody staining cocktail, containing the following antibodies diluted in FACS buffer: CD3-FITC (Biolegend, catalog no. 317306), CD4-APC-Cy7 (Biolegend, catalog no. 357416), CD8a-Alexa Fluor 700 (BioLegend, catalog no. 344724), CD137-BV785 (BD Biosciences, catalog no. 744397), PD-1-BV605 (BioLegend, catalog no. 329924), CXCR5-Alexa Fluor 647 (BD Biosciences, catalog no. 558113), ICOS-PE (BioLegend, catalog no. 313520r), CCR7-BV710 (BioLegend, catalog no. 353228), CD45RA-BV510 (BioLegend, catalog no. 304142). Cells were then washed in FACS buffer and incubated in 100 µl live-dead marker 7-aminoactinomycin D (7AAD) for 10 minutes at 4°C in the dark. For flow cytometric measurements and index cell sorting, cells were resuspended in FACS buffer at 2×10^6^ cells per ml. Single CD8^+^7AAD^−^CD3^+^CCR7^−^ T cells with an activated phenotype based on surface expression of 4-1BB (CD137), PD-1, ICOS or CXCR5 were isolated from freshly thawed PBMC samples or after 10-day *in vitro* expansion culture based on a 7AAD^−^CD3^+^CD4^+^ or 7AAD^−^CD3^+^CD8^+^ phenotype and high expression of CD25 and OX40 or CD69 and CD137, respectively. Cells were analyzed or isolated using a FACS Aria III (BD). Cells were sorted into 384-well plates using the indexed sort option of the Diva software.

### cDNA synthesis and *TR* gene amplification

cDNA synthesis and *TCR* gene amplification were performed as previously described (Wahl et al., 2022a). In brief, cDNA synthesis was performed in the original sort plates using random hexamer primers. *TRA* and *TRB* gene amplification was performed by individual semi-nested PCR using separate *TRAV*- and *TRBV*-specific primer sets. Row- and column-specific barcodes attached to the second PCR primers allowed pooling of all amplicons for next generation sequencing using Illumina MiSeq or NextSeq 2×300.

### TCR sequence analysis

TCR gene sequence analysis and segment annotation were performed using SciReptor (Imkeller et al., 2016; Wahl et al., 2022a). In brief, paired sequencing reads were assembled using PandaSeq setting the maximal and minimal length thresholds to 550 and 300 bp, respectively (Masella et al., 2012), with a minimal overlap of 50 bp and read quality score above 0.8. Reads are assigned to their plate position according to the column- and row-specific barcodes. V, D, and J segments as well as CDR and framework regions (FWR) were identified by Ig BLAST and annotated (Ye et al., 2013). Cell phenotype data from index sort fcs files and metadata were linked to the sequence information and stored in a relational MySQL database. Further analysis of the clonal composition and sequence features were performed in R studio using the following packages: ggplot2, ggalluvial, pheatmap, vegan, stringr, dplyr. Clustering of TCR according to sequence similarity was performed using the BLscore package and further visualization was executed in Cytoscape.

### TCR expression cloning

TCRs were selected for functional characterization based on a scoring system including enriched V segment usage, V segment pairing, clone size, and TCR clustering based on BLscore algorithm (Wahl et al., 2022a). TCRs that scored in one or several categories were cloned. The cloning and expression of TCRs in TCR^neg^CD3^+^CD4^+^ or TCR^neg^CD3^+^CD8^+^ Jurkat cells (J76-CD4, J76-CD8) was performed as previously described (Wahl et al., 2022a). In brief, full-length *TRA* and *TRB* genes were cloned into retroviral pMSCV-PlmC expression vectors. If two productive *TRA* genes were detected in one cell, the one with the higher copy number was chosen for cloning. After cloning and sequence verification, Phoenix Ampho cells (American Type Culture Collection, catalog no. CRL-3213; RRID:CVCL_H716) were transfected with the TCR expression vectors using 2.5 M CaCl_2_ and HEPES buffered saline (HBS) and cultured in Dulbecco’s modified Eagle’s medium (DMEM) GlutaMAX medium (Life Technologies) with 10% heat-inactivated FBS at 37°C and 5% CO_2_. Retroviral particles were harvested from the supernatants the next day and used for the transduction of J76-CD4^+^ and J76-CD8^+^ T cells. For spin infections, 5×10^5^ J76 cells were resuspended in 1 ml retroviral supernatant containing 10ug/ml protamine sulfate, plated in 24-well tissue culture plates, and spun at 2000g and 32°C for 1.5 h. Afterwards, 1ml of RPMI 1640 supplemented with 10% heat-inactivated FBS and 2mM Glutamine was added and cells incubated for 2 days at 37°C and 5% CO_2_. After 7 days of puromycin dihydrochloride (0.8 µg/ml) selection, TCR expression was confirmed by flow cytometric analysis by anti-TCR (Biolegend, catalog no. 306720) and LIVE/DEAD fixable Near-IR dead cell (Invitrogen, catalog no. L34975) staining. Only cell lines with TCR expression levels above 10% TCR^+^ cells were screened for reactivity.

### TCR reactivity testing

TCR reactivity tests and EBV immortalization were performed as described previously (Wahl et al., 2022a). In brief, 2.13 × 10^5^ EBV-immortalized B cells were seeded in 100 µl AIM-V medium into 96-well U bottom tissue culture plate and loaded with peptide (2.5 µg/ml) for 2-4h at 37°C and 5% CO_2_ before addition of 4.27× 10^5^ TCR-transgenic J76 T cells in 100 µl AIM-V medium. For HLA blocking experiments, HLA-DR (Biolegend, catalog no. 307602), HLA-DQ (Abeomics, catalog no. 10-4134), HLA-DP (Leinco Technologies, catalog no. H260), HLA-A (Abeomics, catalog no. 36-2474), HLA-B (Abeomics, catalog no. 36-2475), or HLA-C (Biolegend, catalog no. 373302) blocking antibody was added at a concentration of 10 µg/ml together with T cells (Cassotta et al., 2020). After 24h incubation at 37°C, supernatants were harvested and serially diluted for IL-2 concentration ELISAs in 384-well plates using one quarter of the recommended reaction volumes for the human IL-2 ELISA MAX Deluxe kit (Biolegend). OD values were measured at 450nm with a M1000 Pro plate reader (Tecan). IL-2 concentrations were determined and plotted using Excel and GraphPadPrism 9.

### HLA typing

For HLA typing, gDNA was extracted from PBMCs using the DNeasy Blood and Tissue Kit (Qiagen) according to the manufacturer’s instructions. HLA typing was performed at the Institute for Immunology and Genetics, Kaiserslautern, Germany, or at the DKMS facility, Dresden, Germany.

### Peptide MHC *in vitro* binding assay

PfCSP peptides covering the major predicted binders identified by NetMHCpan4.0 were loaded onto five MHC monomers (Immudex) and coupled to streptavidin beads (Spherotech) following the instruction provided by Immudex. The level of peptide binding was quantified using a fluorescently labelled antibody against a conformational epitope within the ß2-microglobulin domain, which is only accessible when peptide is bound. For each allele, a positive control peptide provided by Immudex and an empty molecule were included as negative controls. Peptide binding was calculated relative to allele-specific negative control as fold change in median fluorescence intensity (MFI) and threshold for binding was set to MFI fold change of 2.

### MHC binding predictions

MHC peptide binding predictions were performed using the NetMHCpan4.0 algorithm (Reynisson et al., 2020). All 8mer, 9mer and 10mer peptides covering the FMP013 immunogen sequence (aa20 −380) were included in the analysis and evaluated for binding to HLA-A and HLA-B alleles expressed by at least one of five donors (F1-F5) enrolled in the FMP013 study. All predicted binders (EL rank <7) were plotted.

